# RNA sequencing reveals the developmental onset of autosomal gene expression differences in male and female extravillous trophoblasts

**DOI:** 10.1101/2024.12.16.628789

**Authors:** Matthew J. Shannon, Alexander G. Beristain

## Abstract

**Background:** The human placenta is an essential organ for fetal development and pregnancy success. Across gestation, blastocyst-derived trophoblasts facilitate the major functions of the placenta. Specifically, invasive trophoblast subtypes, called extravillous trophoblasts (EVT), play central roles in coordinating nutrient accessibility and maternal immunomodulation to semi-allogeneic fetal and placental tissues. Like the fetus, these trophoblasts can be chromosomally male (XY) or female (XX). While male and female trophoblasts are associated with distinct placental gene signatures, specific differences between male and female EVT have not been defined across the first trimester.

**Methods:** To understand how male and female EVT differ, we subjected male and female first trimester EVT cell preparations to bulk RNA sequencing. Concurrently, publicly available single-cell RNA sequencing datasets of first trimester placental and decidual tissues were utilized to resolve EVT differentiation and EVT subtype-specific sex differences. Candidate genes were then selected and immuno-localized to specific regions and cell populations in male and female placentas.

**Results:** We found that before week 10 of gestation, both male and female EVT lineage cells increase expression of transcripts associated with cell proliferation. Sex-related gene differences within this early developmental time-point are restricted to genes residing on sex chromosomes. Following week 10 of gestation, there is a broad up-regulation of genes linked to immunoregulation in male and female EVT. However, within this later developmental period, autosomal gene differences appear in relation to biological sex. We go on to show that these sex-dependent autosomal gene differences influence EVT-maternal cell signalling within the uterus whereby pregnancies exposed to a male placenta demonstrate more complex MIF and CD99 as well as angiogenesis-associated VEGF cell-cell signals between male EVT and the female maternal immune and non-immune cells found throughout the uterus.

**Conclusions:** These findings resolve early first trimester EVT lineage trophoblast sex differences and highlight a developmental timepoint that is critical to male and female autosomal gene expression.

**HIGHLIGHTS:**

- First trimester male and female placenta cells were compared using bulk and single-cell transcriptomics
- Gestational age and placental sex are the leading drivers of variation in gene expression
- Sex-related differences in autosomal genes generally arise on week 10 of gestation
- Sex-related differences in trophoblast and uterine cell crosstalk are driven by autosomal gene differences arising after week 10 of gestation

**PLAIN ENGLISH SUMMARY:** The human placenta is a temporary organ that forms during pregnancy. Importantly, the placenta acts as a surrogate for not yet functioning fetal organ systems (i.e., the heart, lungs, and kidneys) while they mature within the fetus. Because the placenta develops from cells of the early embryo, it can be biologically male or female. Male and female placentas are genetically different from each other, where these differences may differentially influence placental functions and pregnancy health through unknown mechanisms. Therefore, this study aimed to understand how male and female placenta cells differ at the level of gene expression. We find that on week 10 of gestation, but not before, female placenta cells express a different repertoire of genes compared to male placenta cells. We show that these differences potentially affect how cells of the placenta and uterus interact with each other. Taken together, these results identify a developmental timepoint in pregnancy where the biological sex of the placenta may instruct subtle differences in how male or female placenta cells communicate with the maternal compartment in pregnancy.

**Summary Statement:** Bulk and single-cell RNA sequencing provides comprehensive comparison between uncultured male and female HLA-G-purified trophoblasts derived from first-trimester human placentas.

## BACKGROUND

In eutherian mammals, the placenta is the first fully functioning organ of the conceptus to develop. In humans, the placenta rapidly grows and forms the key cellular components required for its function by the end of the first trimester of pregnancy [1,2]. Serving many important functions, the placenta is essential for fetal growth and survival. For example, the placenta anchors the growing fetus to the uterus, coordinates the exchange of nutrients, waste, and oxygen between fetal and maternal compartments, and modifies the maternal immune response to fetal and placental tissues [3–5]. It is generally accepted that alterations in placentation contribute to the development of many pregnancy-related disorders that directly impact the health of the developing fetus and mother.

While the placenta is composed of cells derived from the extra-embryonic mesoderm and trophectoderm, trophectoderm-derived trophoblasts perform the majority of the placenta’s functions. Trophoblasts develop along one of two cell differentiation trajectories, termed villous and extravillous (Fig. 1A) [1,2]. During villous differentiation, progenitor cytotrophoblasts (CTB) undergo asymmetrical cell division and cell fusion to form a multi-nucleated cellular layer called the syncytiotrophoblast (SCT) (Fig. 1B) [6]. The SCT layer serves as the placenta’s endocrine engine and forms the selective barrier between fetal and maternal circulations that promotes the transfer of nutrients and oxygen towards the fetus and minimizes fetal exposure to harmful substances [7,8]. Alternatively, progenitor CTB within areas of chorionic villi that are attached to maternal decidua undergo differentiation along the extravillous pathway [9]. Extravillous differentiation establishes cellular structures called anchoring columns that physically attach the placenta to the maternal compartment. Proliferative trophoblasts within anchoring columns, termed column CTB (cCTB), mature into invasive extravillous trophoblasts (EVT) (Fig. 1B) [1,9]. EVT are involved in important vascular remodelling processes within the uterus and likely modulate the maternal immune response to pregnancy through direct interactions with maternal immune cells [10–12]. EVT-maternal interactions represent intimate self-none-self cell communication that necessitate robust tolerogenic mechanisms to support placenta development and pregnancy health.

**Figure 1:**
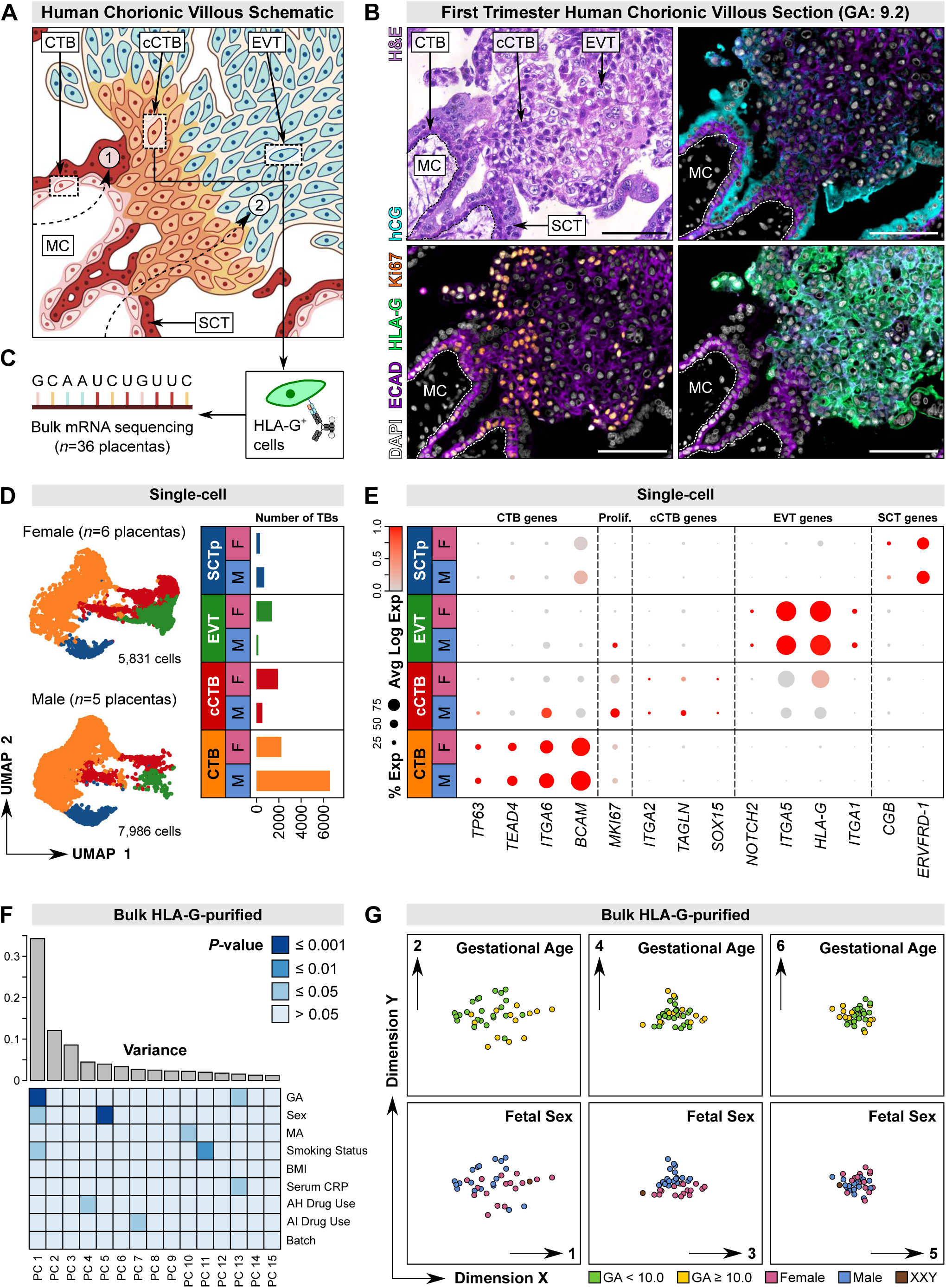
Bulk RNA-sequencing of HLA-G^+^ cells to study first trimester extravillous trophoblast gene expression. (A) Schematic of a first trimester chorionic villous section including anatomical representations of floating and anchoring placental villi. Highlighted are the following trophoblast subtypes: cytotrophoblast (CTB), column cytotrophoblast (cCTB), extravillous trophoblast (EVT), and syncytiotrophoblast (SCT), as well as the villous mesenchymal core (MC). (B) Representative images of serial sections of a first trimester chorionic villus (week 9.2 of gestation) stained with hematoxylin and eosin or combinations of antibodies targeting ECAD, hCG, Ki67, and HLA-G; nuclei are stained with DAPI. Scale bars = 100 μm. Dotted lines separate CTB from the MC. CTB, cCTB, EVT, and SCT regions are indicated by arrows. (C) Schematic summarizing the magnetic immuno-bead cell isolation approach utilized for the purification and bulk RNA-sequencing of first trimester HLA-G^+^ trophoblasts. (D) Left: Uniform manifold approximation and projection (UMAP) plot of 5,831 female (top) and 7,986 male (bottom) first-trimester trophoblasts. Shown are 4 clusters of CTB, cCTB, EVT, and SCTp. Right: Bar plot summarizing the number of female and male trophoblasts within each trophoblast cluster. (E) Dot plot showing the percent and average Log expression level (0.0 to 1.0) of CTB, proliferative (Prolif.), cCTB, EVT, and SCTp gene markers in each cluster split by sex. (F) Top: Bar plot demonstrating the percent of variance explained by the top 15 principal components (PCs) in the bulk RNA-seq data. Bottom: Heatmap correlating each PC with the sample metadata provided in Table S1. (G) Multidimensional scaling plots demonstrating sample clustering across PC1 vs. PC2, PC3 vs. PC4, and PC5 vs. PC6. Samples are colour coordinated based on gestational age (top) or placenta sex (bottom).

Trophoblasts are usually either chromosomally male (XY) or female (XX). Previous studies utilising bulk and single cell transcriptomics have highlighted sex-related differences between male and female trophoblasts, and relatedly, transcriptomic-inferred differences in maternal-placental cell cross-talk [13–18]. However, these studies did not focus on the EVT-specific contributions to these sex differences. As a result, sex differences between male and female EVT invasion, EVT remodeling of maternal vessels, and EVT cross-talk with maternal immune cells remain unknown. Importantly, shallow EVT invasion and inadequate vascular remodelling are associated with obstetrical complications presenting with sex bias, like preeclampsia [19,20]. Because pregnancies carrying a female fetus are at increased risk for preeclampsia within the first trimester [21], it is possible that biological differences between first trimester male and female EVT establish predispositions to specific pregnancy outcomes. To our knowledge, no study has specifically compared male and female EVT at the transcriptomic level before week 10 of gestation. Therefore, a comprehensive comparison of first trimester male and female trophoblasts is needed to determine if EVT-related gene expression is impacted by biological sex.

To address this, we isolated cells of the EVT sub-lineage from human male and female first trimester placentas and subjected these cells to bulk RNA sequencing (RNA-seq). In tandem, we also leveraged single-cell transcriptomic datasets of first trimester placentas and decidual tissues [22,23] to investigate if sex differences in cellular heterogeneity exist along the EVT pathway. Our work suggests that EVT are the most sexually divergent trophoblast cell type in the human placenta. We also show that gestational age and trophoblast sex are the primary drivers of variation in autosomal genes expressed in cells differentiating along the EVT pathway. Specifically, we show that sex-related differences in autosomal gene expression initiate only after 10 weeks’ gestation, and that these differences may result in alterations in how male and female EVT interact with maternal decidual cells. Overall, this work identifies EVT sex-related differences in gene expression that are coordinated in part by developmental processes in the first trimester of pregnancy.

## METHODS

### Patient recruitment and tissue collection

Placental tissues were obtained with approval from the Research Ethics Board on the use of human subjects, University of British Columbia (H13-00640). All samples were collected from pregnant individuals (19 to 39 years of age) providing written informed consent undergoing elective termination of pregnancy at British Columbia’s Women’s Hospital, Vancouver, Canada. First-trimester placental tissues (n=39) were collected from participating patients (gestational ages ranging from 5–12 weeks) having confirmed viable pregnancies by ultrasound-measured fetal heartbeat. Patient clinical characteristics (i.e., height and weight) were obtained to calculate body mass index (kg/m^2^) and all consenting patients provided self-reported information, via questionnaire, to having first-hand exposure to cigarette smoke or having taken anti-inflammation or anti-hypertensive medications during pregnancy. Patient and sample clinical characteristics are summarized in Table S1A.

### Tissue sex determination

Genomic DNA was extracted and purified from all placental tissues with the DNeasy Blood & Tissue kit (Qiagen) using manufacturer’s protocol. DNA purity was confirmed using a NanoDrop Spectrophotometer (Thermo Scientific). 100ng gDNA from each sample was dedicated to polymerase chain reaction using Taq DNA polymerase and combinations of the following forward and reverse primers for the X-chromosome linked transcript ZFX and the Y-chromosome linked transcript ZFY (Chromosome X and Y F: 5’-ATTTGTTCTAAGTCGCCATATTCTCT-3’; Chromosome X R: 5’-GAACACACTACT-GAGCAAAATGTATA-3’; Chromosome Y R: 5’-CATCTTTACAAGCTTGTAGACACACT-3’). 20 μl amplification products were directly analyzed on a 2% agarose gel (data not shown). To confirm sample sex, expression levels of the ZFY, ZFX, and XIST genes were considered after bulk RNA-sequencing. This revealed a likely sex chromosome aneuploidy in sample E12 (XXY). Sample E12 was removed prior to all sex-specific comparisons.

### Maternal CRP Quantification

Maternal serum was collected from all samples processed for bulk RNA-seq (n=36). CRP concentration in maternal serum was measured using the Human CRP rapid ELISA Kit (Invitrogen) according to manufacturer’s instructions. Maternal CRP levels are reported in Table S1.

### HLA-G^+^ trophoblast purification and RNA isolation

Placental chorionic villi were mechanically and enzymatically digested into single cell suspensions, and these suspensions were subjected to density gradient centrifugation followed by magnetic immuno-bead isolation of HLA-G^+^ cells. Briefly, placental villi were finely minced using a razor blade and following this digested twice with 0.125% trypsin (Sigma-Aldrich, St. Louis, MO, USA; T8003) and 0.1 mg/ml DNAse I (Sigma-Aldrich) for 30 min. After digestion, supernatant was collected, centrifuged, and cell pellets were washed in DMEM/F12 containing 10% fetal bovine serum. The resulting cell suspensions were then passed through a 100 μm filter before layering on top of a discontinuous Percoll gradient (10–70%) and centrifugation at 1000 g for 30 min without engagement of the rotor brake. Cell bands residing between the 35% and 50% layers were collected and washed in 1X HBSS.

From the collected cell suspension, HLA-G^+^ cells were isolated using the Miltenyi MS Column microbead positive cell purification kit following manufacturer’s instructions. Specifically, 4 x 10^6^ cells were incubated at 4°C for 30 min with anti-HLA-G-PE (MEM-G/9, 1:100; ExBio, Vestec, Czech Republic) antibody in PBS containing 0.5% BSA and 2 mM EDTA. Following this, cells were incubated with 20 μl anti-PE microbeads (130-048-801, Miltenyi) for every 1 x 10^7^ cells. Finally, HLA-G^+^ cells were collected via positive selection using Miltenyi MS MACS separator columns. Total RNA was then prepared from the HLA-G^+^ cells using TRIzol Reagent (Invitrogen) followed by RNeasy Mini kit (Qiagen) clean-up following the manufacturer’s protocol. RNA purity was assessed using a NanoDrop Spectrophotometer (Thermo Scientific) and Agilent 2100 Bioanalyzer (Agilent Technologies, Santa Clara, CA) prior to bulk RNA-sequencing. Only RNA samples having an RNA Integrity Number (RIN) greater than 7.0 were used for bulk RNA-sequencing.

### Bulk HLA-G^+^ RNA-seq library construction

HLA-G^+^ trophoblast RNA samples (n=36) were sequenced using an Illumina HiSeq 2500 instrument generating 20 million paired reads. Libraries were prepared using the TruSeq Stranded mRNA Library Prep Kit (Illumina, San Diego, CA) and loaded onto the nanofluidic NeoPrep instrument for Poly-A mRNA selection, fragmentation, cDNA synthesis, sequencer adaptor ligation, and amplification. Raw BAM files were then downloaded, mapped against the hg19 reference genome using STAR read aligner, and FPKM read counts were generated using Cufflinks 2. Sample sequencing metrics are summarized in Table S1B.

### Single-cell RNA-seq data acquisition

Droplet-based first-trimester scRNA-seq decidual and placental tissue data (n=12) was obtained from the public Gene Expression Omnibus (GEO) repository (GEO174481) [24] and from the ArrayExpress public repository (E-MTAB-6701) [23]. Raw BAM files from the GEO and ArrayExpress repositories were downloaded and pre-processed as previously described [22,24] using 10X Genomics Cell Ranger version 3.0 software with STAR read alignment and barcode and UMI counting against the hg19 reference genome. Sample sequencing metrics are summarized in Table S1B.

### Bulk RNA-seq data analysis

The code used for bulk RNA-seq data processing and all analyses is available at: https://github.com/MatthewJShannon.

Data pre-processing and quality control: Raw count data from each HLA-G^+^ trophoblast sample was combined, Log_2_ transformed, and filtered to remove genes containing fewer than 1 count in less than 3 samples. Variation due to sequencing batch was removed using an empirical Bayes approach via the “ComBat” function of the SVA package (version 3.38.0) in R.

Principal component analysis: Principal component analysis was conducted on the normalized and filtered data using the “princomp” function of the stats package (version 4.0.5) and visualized using the ggplot2 package (version 3.5.0) in R.

Multidimensional scaling: Samples were mapped in 2-dimensions using the “plotMDS” function of the limma package (version 3.46.0) in R using the normalized and filtered data. During plotting, samples were colour-coded based on the variable of interest.

Heatmaps: Sample-sample Pearson’s correlations were calculated and visualized as a heatmap and dendrogram. To accomplish this, a covariance matrix of the normalized and filtered data was generated and scaled using the “cor” function of the stats package (version 4.0.5). The heatmap and dendrograms were then generated using the “pheatmap” function of the pheatmap package (version 1.0.12) in R. During plotting, a ward.D2 clustering algorithm was selected and cluster distances were calculated using a Euclidean approach. Next, the top 50 most variably expressed transcripts were identified in the normalized and filtered dataset. With this list of genes, another heatmap and dendrogram were generated using the pheatmap package in R. During plotting, gene rows and sample columns were clustered without bias using default parameters by setting “cluster_cols = TRUE” and “cluster_rows = TRUE”.

Gene expression plotting: Gene expression levels were plotted using GraphPad Prism (version 9.3.1).

Differential gene expression: Differential gene expression analyses were performed on raw count data using the DESeq2 package (version 1.30.1) in R. Pairwise comparisons for differential gene expression analyses were based on experimental designs considering either gestational age or sex in combination with the batch variable to control for changes in measured counts related to batch. For each differential gene expression analysis, results were ordered, and transcripts were considered significant at a Benjamini-Hochberg corrected p-value <0.05. Results were visualized using the R package EnhancedVolcano (version 1.8.0) with a p-value cut-off of 10e-5 and for gestational age analyses, a Log_2_ fold change cut-off of 1.00 used as thresholds for significance in data visualization. Comprehensive differential gene expression results are summarized in Table S4.

Gene Ontology: Gene ontology enrichment analysis was performed on the results of our gestational age differential gene expression analysis (i.e., transcripts with a Benjamini-Hochberg corrected p-value < 0.05) using the clusterProfiler package (version 3.18.1) in R. Gene ontology terms pertaining to the “Biological Processes” ontology were considered significant at a Benjamini-Hochberg corrected p-value < 0.05 for genes found enriched in our <9 week gestational age category or our ≥10 week gestational age category. All results were visualized using the “dotplot” function of the ggplot2 package (version 3.5.0) in R.

### Single-cell RNA-seq data analysis

The code used for single-cell RNA-seq data processing and all analyses is available at: https://github.com/MatthewJShannon.

#### Data pre-processing and quality control

To ensure only high-quality cells were used in downstream analyses, cells containing fewer than 500 detected genes and greater than 20% mitochondrial DNA content were removed using the Seurat R package (version 4.0.1) [25]. Individual samples were scored based on expression of G_2_/M and S phase cell cycle gene markers, scaled, normalized, and contaminating cell doublets were removed using the DoubletFinder package [26] (version 2.0.3) with default parameters and a doublet formation rate estimated from the number of captured cells. To minimize bias due to read dropouts the “FindVariableFeatures” function was used with a “vst” selection method to prioritize highly variable genes in each sample for all downstream analyses. Following pre-processing, all samples were merged and integrated using cell pairwise correspondences between single cells across sequencing batches. During integration, the Pearson residuals obtained from the default regularized negative binomial model were re-computed and scaled to remove latent variation in percent mitochondrial DNA content as well as latent variation resulting from the difference between the G_2_/M and S phase cell cycle scores.

#### Maternal-placental interface cell clustering and identification

Following data pre-processing, a total of 64,248 single cells from n=12 placental and decidual samples remained (23,753 cells from n=6 female placentas and 40,495 cells from n=5 male placentas). All cells were clustered in Seurat at a resolution of 0.900 using 30 principal components. Cell identity was determined by observation of known gene markers of the maternal-placental interface (i.e., CD34, KIT, ENTPD1, NCAM1, ITGAX, PTPRC, KLRB1, CD8A, IL7R, CD3G, FCER1A, CD1C, CLEC9A, S100A12, IL1B, FCGR3A, CD14, CD4, LYVE1, VCAM1, COL6A2, DLK1, PDGFRB, MCAM, ACTA2, MYH11, DKK1, IGFBP1, VIM, CD59, TFAP2A, TFAP2C, GATA2, GATA3, TP63, TEAD4, ITGA6, BCAM, MKI67, ITGA2, TAGLN, NOTCH2, ITGA5, HLA-G, ITGA1, CGB, ERVFRD-1) in each cluster. Prior to trophoblast identification, individual states were binned into CTB, cCTB, EVT, SCTp, Hofbauer, uterine natural killer cell (NK), conventional NK, T cell, macrophage, endothelial cell, epiglandular cell, or decidual stromal cell types. Cell projections were visualized using Uniform Manifold Approximation and Projection (UMAP) plots and where relevant, split by sex.

#### Male and female trophoblast sub-setting, cell clustering, and identification

Trophoblasts were discerned from non-trophoblast cells based on the following criteria: First, trophoblasts were selected if they expressed combinations of KRT7, EGFR, TEAD4, TP63, TFAP2A, TFAP2C, GATA2, or GATA3, if they expressed EVT-enriched HLA-G [27], or if they expressed SCT-enriched ERVFRD-1 [28] transcripts at a level greater than zero. Second, trophoblasts needed to express non-trophoblast-related transcripts (VIM, WARS, PTPRC, DCN, CD34, CD14, CD86, CD163, NKG7, KLRD1, HLA-DPA1, HLA-DPB1, HLA-DRA, HLA-DRB1, HLA-DRB5, and HLA-DQA1) at a level equal to zero. After identification, trophoblasts from n=11 placental samples were subset and re-clustered in Seurat at a resolution of 0.375 using 30 principal components. Cell identity was determined as previously described and verified by observation of CTB (TP63, TEAD4, ITGA6, BCAM), proliferative (MKI67), cCTB (ITGA2, TAGLN, SOX15), EVT (NOTCH2, ITGA5, HLA-G, ITGA1), and SCT (CGB, ERVFRD-1) gene markers in each cluster. For all reported single-cell analyses, individual trophoblast states were binned into CTB, cCTB, EVT, and SCTp trophoblast subtypes. Trophoblast projections were visualized using UMAP plots and where relevant, split by sex.

#### Male and female trophoblast-decidual cell clustering and identification

Finally, cells originating from the decidua samples (8,019 cells from n=1 female decidual tissue and 22,669 cells from n=2 male decidual tissues were subset from the male and female maternal-placental interface dataset and merged with the male and female trophoblast datasets. All cells were clustered in Seurat at a resolution of 0.350 using 20 principal components. Cell identity was determined as described previously for the maternal-placental interface object. Prior to trophoblast identification, individual states were binned into CTB, cCTB, EVT, SCTp, macrophage, uterine NK, conventional NK, epiglandular cell, decidual stromal cell, or endothelial cell types. Cell projections were visualized using Uniform Manifold Approximation and Projection (UMAP) plots.

#### Pseudo-bulk principal component analysis

Trophoblast cluster expression matrices (i.e., cCTB, EVT) were subset and state specific expression matrices were independently averaged by sample of origin (n=11) using the “AverageExpression” function. cCTB or EVT sample specific variable features were then re-identified before the “runPCA” function was applied to visualize pseudo-bulk cell states in principal component space.

#### Pearson’s correlation coefficient analysis

Trophoblast cluster expression matrices were independently averaged by sex (male CTB, female CTB, male cCTB, female cCTB, male EVT, female EVT, male SCTp, female SCTp) and Log transformed to create sex-specific gene expression data. Pearson correlation coefficients of the relationship between female and male clusters was calculated using the “cor” function from the stats R package (version 4.0.5).

#### Pseudo-bulk gene expression analysis

Trophoblast expression matrices were independently averaged by sample (n=11) to create sample-specific gene expression data. Gene expression levels were then plotted using GraphPad Prism (version 9.3.1).

#### Pseudotime analysis

The Monocle2 R package version 2.18.0 [29,30] was used to explore the differentiation of progenitor CTB into endpoint EVT in the single-cell data. To accomplish this, trophoblast expression matrices from the CTB, cCTB, and EVT clusters were loaded into Monocle2 for semi-supervised single-cell ordering in pseudotime using YAP1, SPINT1, BCAM, and ELF5 expression to denote cells at the origin and HLA-G, PAPPA2, MMP11, and SERPINE2 gene expression to indicate progress towards EVT. Results were visualized using the “plot_cell_trajectory” function and the ggplot2 package (version 3.5.0) in R.

#### CellChat analysis

Sex-dependent cell-cell communication between EVT and decidual cells of the uterus, as well as between trophoblasts, was determined using the CellChat R package version 2.1.2 [31]. First, the male and female trophoblast-decidual cell data objects were down-sampled using Seurat such that a maximum of 500 cells per cluster, per sex were selected. The normalized and down-sampled Seurat data matrices from the male and female EVT and decidual cell objects were then independently loaded into CellChat to create female and male EVT-uterine data objects. The default CellChatDB ligand-receptor interaction database was then selected and all interactions found within the database were considered. Following this, the cell-cell communication network was inferenced using the “subsetData”, “identifyOverExpressedGenes”, “identifyOverExpressedInteractions”, “projectData”, “computeCommunProb”, “filterCommunication”, “subsetCommunication”, “computeCommunProbPathway”, and “aggregateNet” functions. The “mergeCellChat” function was then used to combine female and male EVT-uterine cell-cell communication networks for direct comparison of cell-cell interactions and pathways. The cell-cell communication analyses were then repeated as described here using the male and female trophoblast datasets. Prior to cell-cell communication analysis, the male and female trophoblast data objects were also down-sampled using Seurat such that a maximum of 500 cells per cluster, per sex were used. Cell-cell communication results are summarized in Table S5.

### Microscopy

#### Histological assessment

Male and female human first trimester placentas (n=3) were fixed in 4% paraformaldehyde overnight at 4°C. Tissues were paraffin embedded and sectioned at 6 µm intervals onto glass slides. Placental tissue sections were then stained with hematoxylin and eosin (H&E) according to standard protocol.

#### Immunofluorescence

Serial placental tissue sections to those dedicated to H&E underwent antigen retrieval by heating slides in a microwave in Reveal Decloaker (Biocare Medical). Sections were permeabilized by Triton X-100 for 5 min at RT followed by incubation with sodium borohydride for 5 min at RT. Slides were then blocked in 5% normal goat serum/0.1% saponin for 1 hr at room temperature and incubated with combinations of the indicated primary antibodies overnight at 4°C: rabbit monoclonal anti-ECAD (clone 24E10; 1:1,000; Cell Signaling Technology), mouse monoclonal anti-HLA-G (clone 4H84; 1:100; Exbio), mouse monoclonal anti-Ki67 (clone B56; 1:100; BD Biosciences), mouse monoclonal anti-hCG (clone 5H4-E2; 1:100; Abcam), and rabbit monoclonal anti-CD99 (clone EPR3096; 1:500; Abcam). Following overnight incubation, sections were washed with PBS and incubated with secondary antibodies for 1 hr at RT: polyclonal Alexa Fluor goat anti-rabbit 568 (1:200; Invitrogen) and polyclonal Alexa Fluor goat anti-mouse 488 (1:200; Invitrogen). Autofluorescence was then diminished using the Vector TrueVIEW Autofluorescence Quenching Kit (Vector Laboratories) and sections were mounted using VECTASHIELD® vibrance™ antifade mounting medium with DAPI (BioLynx).

#### Imaging

All slides were imaged using an AxioObserver inverted microscope (Zeiss) with either a 20x Plan-Apochromat/0.80NA or 40x Plan-Apochromat oil/1.4NA objectives (Zeiss). Clinical characteristics of the samples dedicated to imaging are summarized in Table S1A.

### Statistical analyses

Data are reported as median values with standard deviations. All calculations were carried out using GraphPad Prism (version 9.3.1). Prior to all statistical analyses, data normality was confirmed by Shapiro-Wilk test. For multiple comparisons of normally distributed data, an ordinary one-way ANOVA followed by Tukey’s multiple comparison tests were performed. Differences were accepted as significant at p <0.05. For bulk and single-cell RNA-seq statistical analyses, please refer to the relevant single-cell RNA-seq analysis methods sections found in this manuscript.

## RESULTS

### Trophoblasts of the EVT sub-lineage show sex related differences in gene expression

HLA-G expression is confined to anchoring column trophoblasts (i.e., cCTB) and to invasive EVT and therefore serves as a useful cell-surface marker to isolate cells differentiating along the EVT pathway. Capitalizing on this, an HLA-G immuno-magnetic bead approach was applied to isolate HLA-G^+^ trophoblasts from first trimester chorionic villi (n=36). Following DNA sex typing (n=16 XX, n=19 XY, n=1 XXY) cDNA libraries for bulk RNA sequencing were generated (Fig. 1C). Clinical characteristics and sequencing metrics for all bulk RNA-seq samples are summarized in Table S1. However, because HLA-G is broadly expressed (albeit at different levels) in multiple subtypes of trophoblasts undergoing extravillous differentiation [22], we concurrently leveraged published human trophoblast single-cell transcriptomic data (n=6 XX; n=5 XY) [22,23] to address extravillous cell lineage heterogeneity [22,23]. Clinical characteristics and sequencing metrics for all single-cell RNA-seq samples are summarized in Table S1. Following data preprocessing and filtering, 15,402 transcripts were detected in our bulk RNA-seq data with consistent Log_2_ counts measured in all samples and conditions (Fig. S1A). Across all samples, transcriptomic signatures were between 86% and 100% similar with most samples clustering by gestational age (Fig. S1B; Table S2A). Separately, the single-cell data was processed and filtered as described in Shannon et al. (2024) [22]. This provided 5,831 XX and 7,986 XY trophoblasts for single-cell analysis (Fig. 1D). Four trophoblast clusters were resolved representing 4 major cell types: progenitor CTB (2,227 XX, 6591 XY cells), defined by elevated levels of TP63, TEAD4, ITGA6, and BCAM; anchoring column-associated cCTB (1,925 XX, 522 XY cells), defined by heightened levels of ITGA2, TAGLN, SOX15 and low/moderate levels of HLA-G; EVT (1,359 XX, 171 XY cells), defined by elevated levels of NOTCH2, ITGA5, HLA-G, and ITGA1, and mononuclear precursors of the multinucleated syncytiotrophoblast layer (SCTp; 320 XX, 702 XY cells), defined by elevated levels of CGB and ERVFRD-1 (Fig. 1E).

Principal component analyses were then conducted on both bulk and single-cell datasets to associate data variability with sample metadata. Within the bulk RNA-seq data, 34.3% of variation is explained by the first principal component which is attributed to the gestational age, placenta sex, and maternal smoking status variables. 12.1%, 8.6%, and 4.5% of data variation is further explained by principal components 2, 3, and 4, respectively (Fig. 1F). Principal component 5 most strongly correlates with placenta sex and accounts for 3.95% of variability. The remaining principal components (7-35) contribute minimally to variability in our bulk dataset and correlate with additional clinical characteristics indicated in Table S1A and data are summarized in Table S3. These analyses indicate that placenta sex and gestational age are the largest drivers of variation in gene expression in HLA-G^+^ cells. To visualize these trends and further discern the influence of sex from gestational age, samples were plotted in 2-dimensional space using multidimensional scaling plots. Samples predominantly cluster by gestational age along principal components 1 and 2 and by sex along principal components 3 and 4 (Fig. 1G). Further, samples do not cluster by maternal age, maternal body mass index, maternal smoking status, or sequencing batch (Fig. S1C). This trend is less clear in the single-cell data. When treated as pseudo-bulk, the cCTB and EVT states from each patient do not cluster by gestational age (Fig. S1D). This difference may be due to reduced sample power and cell capture limitations inherent to the single-cell platform. Regarding placenta sex, Pearson’s correlation coefficient analysis between single-cell states shows that transcriptionally, male and female CTB are 99.5% similar, male and female cCTB are 95.1% similar, male and female EVT are 93.2% similar, and male and female SCTp are 98.9% similar (Fig. S1E; Table S2B). Overall, the greatest sex differences in gene expression are attributed to HLA-G-expressing cCTB and EVT. Finally, we removed sample E12 (XXY) and plotted male (XY) and female (XX) expression levels of EVT-associated transcripts (Fig. S1F) in both datasets to directly compare our bulk and single-cell data (Fig. S1G). As expected, gene expression patterns are consistent in both datasets. The exception is HLA-G which shows higher levels in female EVT from the single-cell dataset but no sex related difference in the mixed population of HLA-G^+^ cells in the bulk RNA-seq dataset (Fig. S1G).

### Gene expression variation in HLA-G^+^ trophoblasts is driven by gestational age and biological sex

Having undertaken preliminary transcriptomic assessment of the HLA-G^+^ trophoblasts, we next aimed to understand how biological sex correlates with clinical characteristics like gestational age. To accomplish this, we extracted the 50 most variably expressed genes and used this gene signature to conduct an unsupervised hierarchical clustering of all HLA-G^+^ samples (Fig. 2A). Consistent with our bulk RNA-seq principal component analysis, samples first cluster according to gestational age (i.e., into <10 weeks’ or ≥10 weeks’ gestation groups) with expression levels of most of the top 50 high variable genes increasing in HLA-G^+^ cells of the ≥10-week gestational age category (Fig. S2). Our rationale for establishing the gestational age threshold at week 10 of gestation relates to the timing of when intervillous blood supply significantly increases as a result of EVT spiral artery plug erosion; this takes place at or shortly after week 10 of gestation [32,33]. However, unsupervised clustering showed an imperfect segregation of samples in the <10 weeks’ and ≥10 weeks’ gestational age categories. This lack of a clear segregation may be due, in part, to inherent limitations for how gestational age is determined in our clinical cohort. Specifically, gestational age was determined by self-reporting the day of last menstruation. These gestational age estimates can vary by ±1 week [34]. Following gestational age, samples next cluster by biological sex within each gestational age category (Fig. 2A). When additional clinical characteristics like maternal age, body mass index, smoking status, or anti-inflammatory/anti-hypertensive drug use during pregnancy are considered no further trends are observed. Overall, unsupervised clustering supports that gestational age and placental sex are the primary drivers of variation in gene expression.

**Figure 2:**
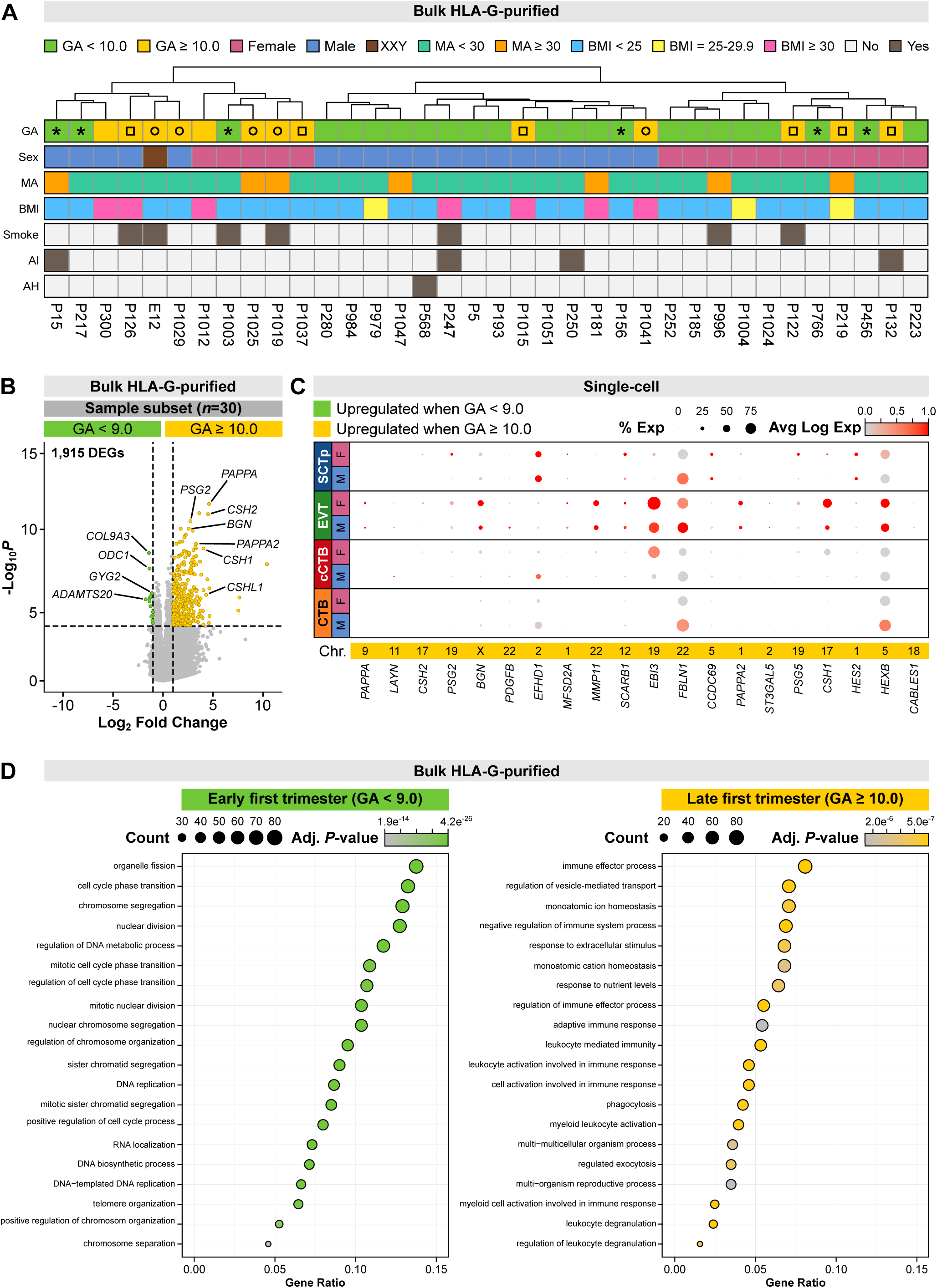
Transcriptomic comparison between early and late first trimester HLA-G^+^ trophoblasts. (A) Dendrogram showing the results of unsupervised hierarchical clustering of all HLA-G^+^ samples, informed by the high variable gene signature presented in Fig. S2. (B) Volcano plot showing differentially expressed transcripts between early first trimester (gestational age <9.0 weeks) and late first trimester (gestational age ≥10.0 weeks) HLA-G^+^ trophoblasts. (C) Dot plot showing the percent and average Log expression level (0.0 to 1.0) of the top 20 adjusted p-value-ranked transcripts identified in Fig. 2B across all single-cell trophoblast clusters split by sex. (D) Top biological processes enriched in the early and late first trimester gene signatures identified in Fig. 2B.

Within the high variable genes, specific Y chromosome transcripts (UTY, RPS4Y1, and DDX3Y) were ranked as the most variable due to their absence in female cells (Fig. S2). Of the remaining transcripts, many have been previously linked to placenta biology, including SPP1 [18] which encodes secreted phosphoprotein 1, the placental growth hormone and placental lactogen encoding CSH1 and CSH2 transcripts [35], and the endocrine-associated CYP19A1, CGA, and CGB5 transcripts [18]. Many transcripts also associate with known EVT functions. These include transcripts that regulate EVT invasion (i.e., KISS1, GDF15, TFPI2, TIMP3) [36–39], transcripts known to correlate with biological functions of EVT (i.e., SERPINE1, PAPPA, PAPPA2, AOC1, PRG2) [40] and transcripts enriched in EVT-differentiated human trophoblast stem cells (i.e., CSHL1, PAEP, GH2, RRAD, SEMA3B, TAC3, DCN, HOPX, BGN) [41,42]. Of the remaining transcripts, 9 belong to the pregnancy specific glycoprotein (PSG) family and another 4 encode extracellular matrix collagens (i.e., COL6A2, COL3A1, COL1A1, COL1A2).

Because gestational age is the leading source of transcript variability, we next set out to examine the impact of gestational age on trophoblast gene expression independent of placental sex. To minimize concerns regarding gestational age determination, six samples with a gestational age equal to week 9 of gestation were removed from our <10 weeks’ gestation category. The remaining samples (gestational age <9 weeks, n=17; gestational age ≥10 weeks, n=13) were subjected to a differential gene expression analysis. 1,915 differentially expressed transcripts were identified between the early and late HLA-G^+^ first trimester trophoblasts; samples aged less than 9 weeks’ gestation had fewer enriched genes and these included COL9A3, ODC1, GYG2, and EVT-associated ADAMTS20 [43] (Fig. 2B; Table S4A). Late first trimester samples show more genes expressed at higher levels than early first trimester samples, including PAPPA, PSG2, CSH2, BGN, ANGPT2, and CSHL1 identified previously as some of the most variable genes (Fig. S2). The top 20 adjusted p-value-ranked transcripts were then extracted and examined across distinct trophoblast subtypes using the single-cell dataset (Fig. 2C). Many of the transcripts are lowly expressed, likely due to differences in mRNA read depth between the bulk and single-cell platforms. However, expression levels of transcripts associated with EVT maturation (i.e., MMP11, PAPPA2, CSH1, EBI3) [40] as well as transcripts not previously studied in EVT biology (i.e., BGN) consistently become enriched towards the EVT single-cell state as well as in the late first trimester HLA-G^+^ samples (Fig. 2C), supporting their role in mature EVT biology. Transcript levels were also split by biological sex. Generally, gene expression levels do not appear to differ between XY- and XX-typed samples (Fig. 2C). However, EBI3 and CSH1 levels are higher in XX cCTB while EFHD1 levels are higher in XY cCTB. As well, FBLN1 levels are higher in XY CTB, SCTp, and EVT while HEXB levels are higher in XY CTB and female SCTp (Fig. 2C).

To examine if distinct biological processes are enriched in EVT lineage cells as a function of gestational age, we next conducted gene ontology analyses on the established early and late first trimester gene signatures (Fig. 2D). Consistent with what we know about EVT maturation, we find that EVT-lineage cells in the early first trimester show enrichment of terms associated with cell proliferation, including the terms ‘organelle fission’, ‘cell cycle phase transition’, and ‘chromosome segregation’ (Fig. 2D). These cell proliferation/mitosis terms likely coincide with the prevalence of highly proliferative HLA-G^+^ comprising the anchoring cell column at this earlier gestational timepoint [44]. After week 10 of gestation, HLA-G^+^ trophoblasts show enrichment of terms associated with immune cell processes and include the terms ‘immune effector process’, ‘negative regulation of immune system process’, and ‘adaptive immune response’ (Fig. 2D). These biological processes are consistent with immunogenic functions of mature EVT [12,27]. Taken together, these data show that even within the first trimester of pregnancy, gestational age differences are associated with profound differences in gene expression in EVT lineage cells.

### Male and female trophoblast autosomal gene differences initiate during week 10 of gestation

Beyond gestational age, we show that the biological sex of the placenta is another leading source of variation in trophoblast gene expression (Figs. 1F, 1G). Single-cell analyses pinpoint cCTB and EVT as the main trophoblast cell types contributing to sex-related gene expression differences in the first trimester human placenta (Fig. S1E). To examine biological sex differences in trophoblasts in more detail, we removed sample E12 (XXY) to conduct a differential gene expression analysis between male (XY) and female (XX) samples (n=35) using our bulk RNA-seq dataset. Independent of gestational age, 26 transcripts were identified as differentially expressed between male and female cells; notably all of these genes mapped to sex chromosomes including the pseudo-autosomal transcript CD99 [45,46] (Fig. S3A; Table S4B). Despite escaping X-chromosome inactivation [47,48] transcript levels of CD99 were surprisingly elevated in male HLA-G^+^ trophoblasts. CD99 encodes a cell surface glycoprotein that, to our knowledge, has not been localized within the first trimester human placenta. Therefore, we probed first trimester XY and XX chorionic villi specimens with CD99 antibody. Using immunofluorescence microscopy, we show a strong CD99 signal in the syncytiotrophoblast and a slightly weaker signal in proximal cells of anchoring columns (Fig. 3A).

**Figure 3:**
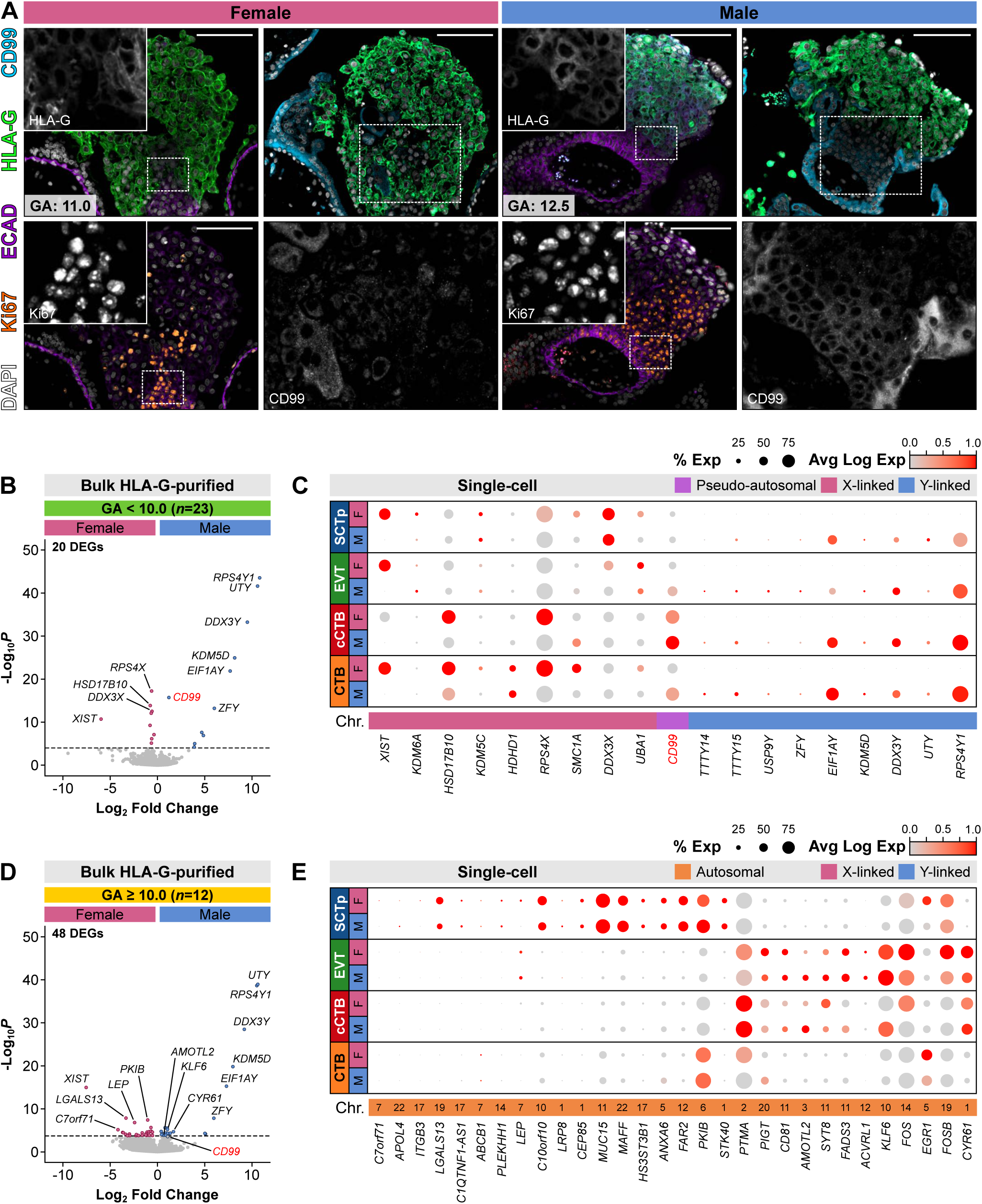
Transcriptomic comparison between first trimester male and female HLA-G^+^ trophoblasts. (A) Representative IF images of serial sections of one late first trimester female (gestational age: 11.0) and one male (gestational age: 12.5) chorionic villus; sections were labeled with antibodies targeting HLA-G, ECAD, Ki67, and CD99. Nuclei are stained with DAPI. Hashed lines depict enlarged insets of HLA-G, Ki67, or CD99 signal. Scale bars = 100 μm. (B) Volcano plot showing differentially expressed transcripts between early first trimester (gestational age <10.0) female and male HLA-G^+^ trophoblasts. (C) Dot plot showing the percent and average Log expression level (0.0 to 1.0) of the significant transcripts identified in Fig. 3B across all single-cell trophoblast clusters split by sex. (D) Volcano plot showing differentially expressed transcripts between late first trimester (gestational age ≥10.0) female and male HLA-G^+^ trophoblasts. (E) Dot plot showing the percent and average Log expression level (0.0 to 1.0) of the significant autosomal transcripts identified in Fig. 3C across all single-cell trophoblast clusters split by sex. Abbreviations are as in Fig. 1.

Considering the influence and possible interaction of gestational age with placenta sex, we next split our male (XY) and female (XX) samples into early (gestational age <10 weeks; n=23) and late (gestational age ≥10 weeks’; n=12) first trimester groups and repeated the differential gene expression analysis. Within the early first trimester group, 20 transcripts were identified as differentially expressed between male and female cells (Fig. 3B; Table S4C). These transcripts were mostly identical to the genes identified when gestational age is not considered and likewise mapped exclusively to the X and Y chromosomes. Genes with increased expression in the early first trimester female HLA-G^+^ cells included the long non-coding XIST transcript, a major effector of X chromosome inactivation [49], as well as RPS4X which encodes ribosomal protein S4, and HSD17B10 which escapes X chromosome inactivation and encodes the 17β-hydroxysteroid dehydrogenase involved in sex steroidogenesis [50]. Genes with increased expression in the early first trimester male HLA-G^+^ cells include the functional RPS4X equivalent RPS4Y1 [51], the histone demethylating transcript UTY [52], and again CD99. All 20 differentially expressed transcripts were then extracted and examined across distinct trophoblast subtypes using the single-cell dataset (Fig. 3C). All X-linked transcripts showed increased expression across all female cells with HDHD1 showing a specific increase in expression within progenitor CTB, HSD17B10 and RPS4X showing increased expression within the CTB and cCTB states, and UBA1 expression being specifically increased within the EVT state. Similarly, all Y-linked transcripts were broadly expressed and specific to male cells in our single-cell dataset. CD99 transcript expression in the single-cell dataset matches what was found by immunofluorescence with elevated expression observed in male column-associated cCTB (Fig. 3C).

When ≥10 week gestation cells are examined, 48 transcripts are differentially expressed between male and female cells (Fig. 3D; Table S4D). Notably, 32 of these genes map to autosomes, including LGALS13, PKIB, LEP, and the RNA gene C7orf71 found with increased expression in female HLA-G^+^ cells as well as KLF6, CYR61, and AMOTL2 found with increased expression in male HLA-G^+^ cells (Fig. 3D; Table S4D). While CD99 was not differentially expressed it does trend towards increased expression in the late first trimester male HLA-G^+^ cells with a p-value of 0.065 (Table S4D). 30 of the 32 autosomal transcripts found differentially expressed in the late first trimester samples were then extracted and examined across distinct trophoblast subtypes using the single-cell dataset (Fig. 3E). LOC101927354 and PSG10P were not represented in the single-cell dataset and were unable to be plotted. The majority of transcripts identified with increased expression in female HLA-G^+^ cells were predominantly expressed by SCTp in the single-cell dataset. However, LEP which was increased in female HLA-G^+^ cells and encodes the hormone leptin was specific to the EVT state (Fig. 3E). Transcripts increased within male HLA-G^+^ cells were more closely associated with cells of the extravillous lineage in the single-cell dataset. For example, AMOTL2, encoding the protein angiomotin-like 2 as well as FADS3, CD81, KLF6, and CYR61 were specifically increased in male cCTB and EVT states (Fig. 3E). However, transcripts like EGR1, FOS, and FOSB show opposing directionality in expression in HLA-G^+^ cells (bulk RNA-seq dataset) and specific cell states (single-cell RNA-seq dataset). For example, where male HLA-G^+^ cells express higher levels of EGR1, encoding a transcription factor previously shown to be enriched in progenitor CTBs [22], female CTB express higher EGR1 levels than male CTB in the single-cell dataset (Fig. 3E).

To understand how the differences in autosomal gene expression that arise in the late first trimester relate to distinct stages of extravillous trophoblast differentiation, we subset progenitor CTB, immature cCTB, and mature EVT states from the single-cell data and performed Monocle2 pseudotime analysis [29,30]. As expected, Monocle2-directed ordering predicts a CTB state origin that develops first into the cCTB then the EVT state (Fig. S3B). Using this trajectory, we next sought to examine how the sex-related differences in gene expression, identified in the bulk RNA-seq data, map across extravillous trophoblast differentiation. We first selected CD99, a transcript with expression increased in early (<10 weeks’ gestation) male HLA-G^+^ trophoblasts for plotting gene expression across differentiation. In both male and female trophoblasts progressing along pseudotime from CTB to EVT, CD99 levels transiently increase as CTB transition into the cCTB state and, following this, decrease to levels observed in CTB as these cells become EVT (Fig. S3C). Notably, and consistent with both bulk and single-cell differential gene expression analyses, the CD99 peak is more pronounced in male cCTB (Fig. S3C). We next selected two transcripts having increased expression in late (≥10 weeks’ gestation) HLA-G^+^ trophoblasts: LEP, which was found increased in female cells and AMOTL2 which was found increased in male cells. LEP levels continue to increase as CTB differentiate into cCTB and EVT, where this stepwise increase across pseudotime is most notably observed in female trophoblasts and is less evident in male trophoblasts (Fig. S3C). While AMOTL2 levels do not increase as drastically as LEP across pseudotime, female cells show a transient increase in AMOTL2 expression in cCTB followed by a subsequent decrease in expression as cCTB progress into EVT; in male cells the increase in expression in cCTB is maintained as cells differentiate into EVT (Fig. S3C).

While we show that these autosomal sex differences initiate after week 10 of gestation, the exact timing of this onset was not clear. Therefore, to map the timing of this onset across the first trimester we categorized our bulk RNA-seq samples into weekly gestational age increments (weeks 6-12 of gestation) and plotted gene expression of the 32 autosomal transcripts identified as differentially expressed between male and female samples across each week of gestation (Fig. S3D). These comparisons are limited by the number of samples in each gestational age sub-category. Nonetheless, we do observe that divergences in the expression of specific genes in male and female HLA-G^+^ trophoblasts initiate on week 10 of gestation, not sooner. Overall, these data show that autosomal gene sex differences generally arise at the later stages of the first trimester in cells progressing along the EVT pathway.

### Sex-related trophoblast gene variances predict cell-cell communication differences in the maternal-placental interface

Because EVT interact with diverse cell types in the uterine environment, it is conceivable that differences in gene expression within male (XY) and female (XX) EVT may lead to modifications in cell-cell communication. To address this, uterine-specific immune and non-immune cell types were re-incorporated into the trophoblast single-cell RNA-seq dataset (Fig. 4A). Following data pre-processing and filtering, we obtained 8,019 decidual cells from one pregnancy (gestationally aged week 10.0) having a female fetus and 22,669 decidual cells from two pregnancies (gestationally aged weeks 9.6 and 12.0) having a male fetus. In addition to the four subtypes of single-cell informed trophoblasts (i.e., CTB, cCTB, EVT, SCTp), we resolve 3 decidual immune cell (i.e., uterine NK, conventional NK, macrophage), one endothelial cell, one epiglandular cell, and one decidual stromal cell cluster(s) (Fig. 4A). Genes functioning as known cell-type markers were used to define cell-specific clusters (Fig. S4A). Examining CD99, along with transcripts encoding the CD99 receptors PILRA and PILRB to cell states within placental and uterine cells showed that CD99 is broadly expressed by most cell types, including trophoblasts and non-trophoblasts (Fig. S4B). CD99 levels, however, are highest in uterine NK, conventional NK, stromal, and endothelial cells. PILRA and PILRB levels, on the other hand, are more specific with PILRA expression highest in macrophages and stromal cells and PILRB expression highest in uterine NK and endothelial cells (Fig. S4B). Expression of LEP and its receptor LEPR is more specific within placental and uterine cells with LEP being specifically expressed in cells of the EVT state and LEPR expression being specific to endothelial cells (Fig. S4B). We next considered AMOTL2 expression alongside MAGI1 and CDH5, two genes encoding proteins previously shown to complex with the angiomotin-like 2 scaffold protein [53,54]. Similar to LEP, expression of AMOTL2, MAGI1, and CDH5 was specific to trophoblasts and endothelial cells (Fig. S4B). Interestingly, we observe differences in gene expression in maternal (female) cells depending on whether placental/trophoblast cells of that pregnancy are male (XY) or female (XX). For example, more cells express CD99 in uterine NK from male pregnancies whereas CD99 expression levels are higher in female pregnancy endothelial cells (Fig. S4B). Inhibitory PILRA levels are higher in macrophages from male pregnancies whereas activating PILRB levels are higher in uterine NK from male pregnancies (Fig. S4B). LEPR, in contrast to LEP levels that are highest in female EVT (Fig. S3C), are expressed by more endothelial cells in male pregnancies (Fig. S4B). While AMTOL2 and CDH5 are expressed by more male cCTB, no differences in AMOTL2, MAGI1, or CDH5 expression levels are observed between pregnancies with a male or female placenta (Fig. S4B). Taken together, ligand and receptor gene expression patterns infer broad CD99 signalling across placental and uterine cells and specific LEP and AMOTL2 communication between trophoblasts and endothelial cells.

**Figure 4:**
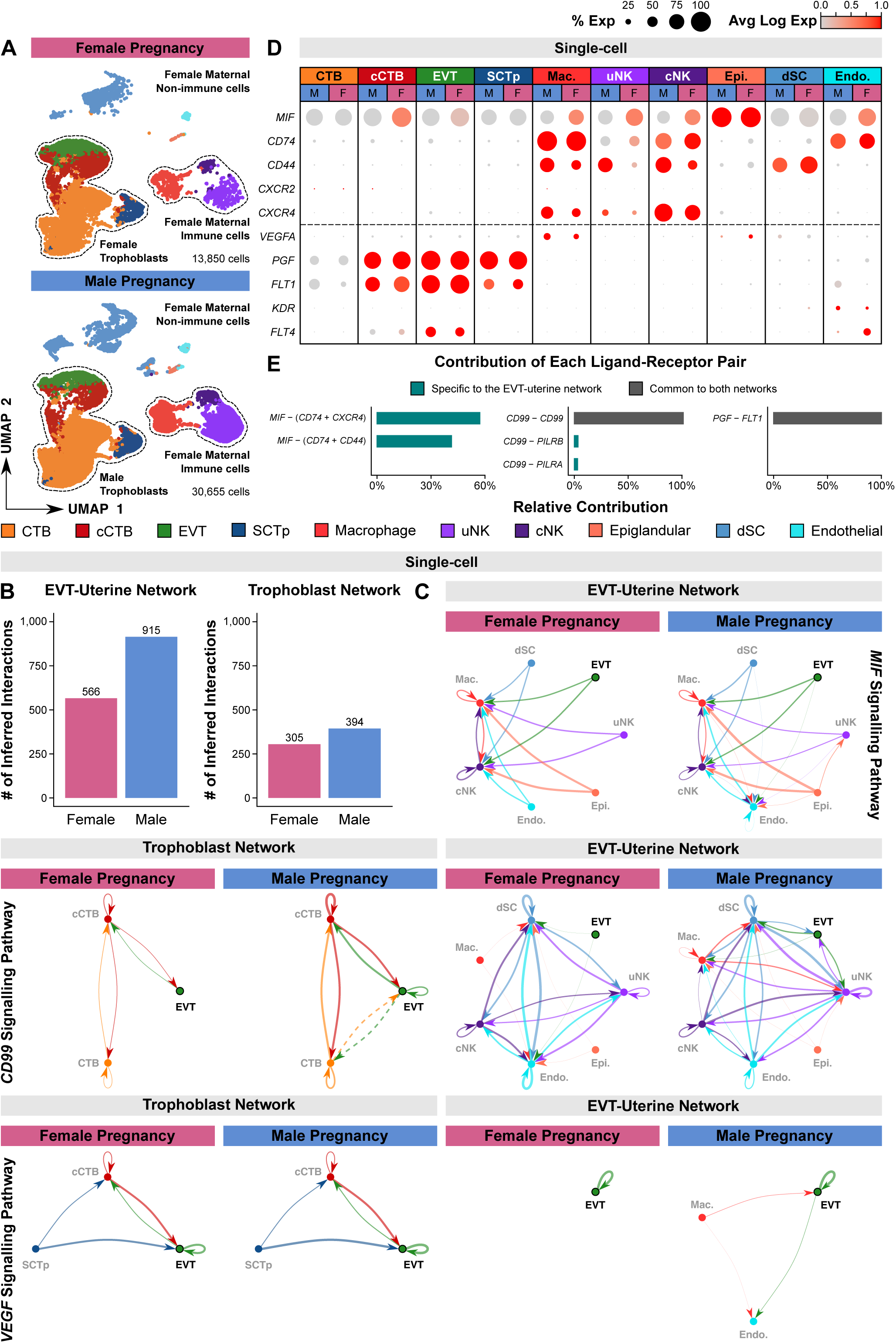
Biological sex-informed cell-cell interactions between trophoblasts and decidual cells. (A) Uniform manifold approximation and projection (UMAP) plot of trophoblasts and decidual cells exposed to either female (top; 13,850 cells) or male (bottom; 30,655 cells) fetal tissue. Shown are 10 clusters of CTB, cCTB, EVT, SCTp, macrophage, uterine natural killer cell (uNK), conventional natural killer cell (cNK), epiglandular cell, decidual stromal cell (dSC), and endothelial cell. (B) Bar plots demonstrating the number of inferred interactions within the female and male EVT-uterine (left; 3,281 cells from the pregnancy with a female placenta and 4,281 cells from the pregnancies with a male placenta) and trophoblast-trophoblast (right; 1,953 female cells and 2,000 male cells) cell communication networks. (C) Circle plots showing cell signalling pathways within the trophoblast-trophoblast (left) and EVT-uterine (right) signalling networks of pregnancies with a female or male placenta. Shown are the inferred MIF (top), CD99 (middle), and VEGF (bottom) signalling networks. Line widths are proportional to the predicted communication probability between the sender and receiver cells. Dashed lines indicate predicted direct CD99-CD99 cell-cell signals that are anatomically unlikely to occur. (D) Dot plot showing the percent and average Log expression level (0.0 to 1.0) of the MIF ligand, the MIF receptors CD74, CD44, CXCR2, and CXCR4, as well as the VEGFA and PGF ligands and the corresponding VEGF signaling pathway receptors FLT1, KDR, and FLT4 in each cell state, split by sex. (E) Bar plots demonstrating the relative contribution of known ligand-receptor pairs to the overall MIF, CD99, and VEGF cell communication networks. Relative contributions are measured as the ratio of the communication probability between each ligand-receptor pair and that of the entire signalling pathway.

To examine if EVT-uterine or trophoblast-trophoblast cell-cell communication is impacted by trophoblast sex, we used CellChat [31] for communication network analysis. To control for differences in cell number between pregnancies with a male or female fetus, a random sample of no more than 500 cells from each cluster was used for all cell-cell communication analyses. This provided 4,281 EVT and decidual cells (EVT-uterine network) as well as 2,000 trophoblasts (trophoblast network) from the pregnancies with a male fetus and 3,281 EVT-uterine network cells as well as 1,953 trophoblast network cells from the pregnancy with a female fetus. 566 significant ligand-receptor interactions are inferred in the EVT-uterine network of the female pregnancy while 915 significant ligand-receptor interactions are inferred in the EVT-uterine network of the male pregnancies (Fig. 4B). Within the inferred interactions of the EVT-uterine network, 70 signalling pathways are enriched with 46 pathways, including CD99, having increased communication probability in cells derived from male pregnancies and 4 pathways having increased communication probability in cells derived from the female pregnancy (Table S5). Within the trophoblast network, 305 and 394 significant ligand-receptor interactions are inferred amongst male and female trophoblasts, respectively (Fig. 4B). Considering the associated signalling pathways, 33 pathways are enriched with 20 pathways, again including CD99, having increased communication probability amongst male trophoblasts and the angiopoietin-like signalling pathway, known to be involved in angiogenesis processes [55], being the only pathway with increased communication probability in female trophoblasts (Table S5).

Within the EVT-uterine cell-cell interactions, the MIF signalling pathway has the greatest sum of inferred communication probabilities in male and female pregnancies (Table S5). MIF ligand-receptor interactions extending from the EVT state predominantly interact with macrophage and conventional NK cell states, however, an extra interaction is observed between male EVT and endothelial cells derived from the pregnancies with a male fetus. This extra interaction is not observed in the pregnancy with a female fetus (Fig. 4C). Examining MIF, and the transcripts encoding the MIF receptors (CD74, CD44, CXCR2, and CXCR4) to cell states using the expanded single-cell dataset shows that MIF is broadly expressed and expressed at higher levels in female cCTB and EVT state cells as well as macrophages, uNK, cNK, and endothelial cells derived from the female pregnancy (Fig. 4D). The MIF receptors CD74 and CD44 are generally similar with specific expression in macrophage, uterine NK, and conventional NK cell states. However, the CD74 receptor is also expressed by endothelial cells while the CD44 receptor is additionally expressed by decidual stromal cells (Fig. 4D). Of the remaining receptors, CXCR2 is lowly expressed by all placental and uterine cells and CXCR4 expression is specific to the macrophage, uterine NK, and conventional NK states (Fig. 4D). Taken together, ligand and receptor gene expression patterns support that EVT-uterine MIF signals predominantly occur between EVT, macrophages, and NK cells found in the decidua. Considering the relative contribution of these known ligand-receptor pairs, we find that MIF interactions are dominated by MIF ligand and the multimeric CD74/CXCR4 or CD74/CD44 receptor complexes (Fig. 4E).

Of the remaining pathways, CD99 signalling demonstrated increased communication probabilities in both the EVT-uterine and trophoblast networks of the pregnancies with a male fetus (Table S5). Considering the CD99 signalling pathway, broad ligand-receptor interactions are seen between EVT and most cells of uterine origin (Fig. 4C), however, more CD99 signalling interactions are observed between cells derived from male pregnancies. While our transcript data highlight male and female expression differences in the cCTB state, CD99 interactions between EVT and uterine cells derived from pregnancies with a male or female fetus do differ. Specifically, CD99 expressed by female EVT interacts only with decidual stromal and endothelial cells in the female pregnancy. CD99 expressed by male EVT, however, interacts with decidual stromal, macrophage, and uterine NK cells in the male pregnancies (Fig. 4C). Amongst trophoblasts, CD99 interactions are generally similar with increased interaction strength and additional EVT-EVT interactions found between male trophoblasts (Fig. 4C). Considering the relative contribution of known ligand-receptor pairs, CD99 signalling predominantly occurs through self-dimerization with CD99. To a lesser extent, CD99 is predicted to interact with the inhibitory PILRA receptor expressed in macrophages and decidual stromal cells and the activating PILRB receptor expressed in uterine NK and endothelial cells of the decidua (Figs. S4B; 4E). Amongst trophoblasts, CD99 interactions are predicted to only occur through CD99 self-dimerization (Fig. 4E). Beyond CD99, the LEP signalling pathway did not meet the thresholds for CellChat significance and was unable to be explored further.

Alternatively, the VEGF signalling pathway is known to regulate vasculogenesis and angiogenesis processes within the human placenta, influence EVT maturation towards an endovascular phenotype, and correlate with the pathogenesis of early onset preeclampsia [56–59]. VEGF ligand-receptor interactions are found to be significant in both the EVT-uterine and trophoblast communication networks (Table S5). Interestingly, while the EVT-uterine network VEGF ligand-receptor interactions are restricted to female EVT-EVT events, more VEGF signals are observed in male pregnancies with ligand-receptor interactions occurring between male EVT-EVT and EVT-uterine macrophage and endothelial cells (Fig. 4C). Within the trophoblast network, no differences are observed between the predicted male and female VEGF ligand-receptor interactions (Fig. 4C). Examining the VEGFA and PGF ligands, as well as the transcripts encoding the VEGF pathway receptors FLT1, KDR, and FLT4 in the expanded single-cell dataset shows that VEGFA expression is highest in macrophages and epiglandular cells while PGF expression is highest in SCTp, cCTB, and EVT (Fig. 4D). While KDR expression is specific to endothelial cells, FLT4 expression is specific to the extravillous-associated cCTB and EVT states as well as endothelial cells found in the decidua (Fig. 4D). We further observe that more endothelial cells express FLT1 in male pregnancies while FLT4 gene expression is higher in decidual endothelial cells derived from the female pregnancy (Fig. 4D). Considering the relative contribution of these ligand-receptor interactions, we find that the VEGF pathway is driven by the PGF ligand and the FLT1 receptor in both the EVT-uterine and trophoblast networks (Fig. 4E).

In summary, we demonstrate that first trimester EVT sex differences in gene expression influence cell-cell communication within trophoblasts and between trophoblasts and diverse uterine cell types. Specifically, transcripts expressed by EVT are predicted to interact with important maternal immune and endothelial cell types in a trophoblast sex-dependant manner.

## DISCUSSION

Using bulk and single-cell transcriptomics, we show that biological sex and developmental timing in early pregnancy together impact the expression of a discreet number of autosomal genes in human trophoblasts. We further provide evidence that biological sex-associated differences in autosomal genes initiate around the end of the first trimester. These differences in gene expression between male and female trophoblasts are largely centered around immuno-modulatory processes within the complex cellular landscape of the uterus. Notably, we provide evidence that biological sex-related gene expression differences occur predominantly in cells committed to the extravillous differentiation trajectory. This observation is interesting as EVT are cells that physically interact with diverse subtypes of maternal cells, suggesting that EVT-maternal cell interactions are in part modulated by biological sex. It is important to note when interpreting this finding that our single-cell dataset lacks true SCT. Therefore, understanding how sex differences within the multinucleated syncytiotrophoblast layer of the placenta compare to those reported here in extravillous-associated trophoblasts is needed.

Supported by both bulk and single-cell transcriptomic evidence, we show that within the first trimester gestational age drives the majority of variation in extravillous lineage trophoblast gene expression. We identify week 10 of gestation as a key developmental timepoint where major differences in gene expression between early and late first trimester cells are observed. This developmental window aligns with the physiological changes that occur alongside the erosion of maternal spiral artery plugs, resulting in an increase in maternal blood delivery to the intervillous space [32,33]. Therefore, it is not surprising that the combined changes in local oxygen tension, exposure to blood-born growth factors and nutrients, increases in fluid shear stress, and the influx of peripheral leukocytes to the intervillous space, all resulting from the erosion of extravillous trophoblast plugs, lead to alterations in the early versus late EVT transcriptome. Our findings suggest that prior to plug erosion (i.e., <10 weeks’ gestation) cells of the EVT lineage harbor mainly pro-mitotic gene signatures. However, after week 10 of gestation and presumably following plug erosion, EVT lineage cells show enrichment in transcripts aligned with immunogenic and matrix remodelling processes. While biological sex differences in autosomal gene expression are not detected prior to week 10 of gestation, we further find that autosomal sex-based gene expression differences in HLA-G^+^ trophoblasts initiate during this physiologically relevant period. While these differences are based on the identification of only 32 differentially expressed autosomal-linked genes, it is possible that these subtle sex-related differences program male and female EVT to differentially respond to, or interact with, their tissue microenvironment. Whether EVT sex-related differences in autosomal genes are further potentiated during adverse pregnancy conditions (e.g., infection, hypoxia) remain to be understood, this work provides insight into baseline male and female differences in EVT biology. To our knowledge, the possibility that biological sex modulates trophoblast gene expression within this distinct developmental window of the first trimester of pregnancy has not been previously reported.

While a major focus of our analyses involved HLA-G^+^ cells isolated from first trimester placental preparations using immune-magnetic HLA-G beads, the heterogeneous composition of this cell population was further resolved using publicly available single-cell datasets. CD99, a pseudo-autosomal gene, was identified as one of the few transcripts showing sex bias in gene expression in HLA-G^+^ cells of the early first trimester. While escaping X-inactivation, CD99 levels were elevated in male trophoblasts. Single-cell transcriptomic analyses further determined that the greatest discrepancy in male versus female CD99 gene expression was in cells of the anchoring column (i.e., cCTB) that are early in progression along extravillous differentiation. CD99 encodes a transmembrane glycoprotein involved in cell surface MHC class I expression [60,61], however, it also plays a role in controlling cell differentiation potential within tumorigenic settings [62–64] and facilitates transendothelial leukocyte trafficking [65,66]. Cell-cell interaction modeling suggests that CD99 may play a role in female uterine immune cell interactions with male or female EVT. However, the significance of sex-biased CD99 expression within cCTB found in anchoring columns is less clear and warrants further investigation. Cell-cell modeling also predicts strong CD99-faciliated interactions between progenitor CTB populations, cCTB, and EVT. This suggests that CD99 may control differentiation of select CTB progenitors committed to the EVT pathway. In addition to CD99, cell-cell interaction modeling suggests that the MIF and VEGF pathways play important roles in EVT-immune cell interactions. MIF encodes a multifunctional cytokine that is abundantly present within the first trimester human placenta [67]. MIF promotes EVT migration and cell invasion, promotes endovascular EVT maturation, and promotes trophoblast as well as decidual cell survival [67–70]. Importantly, an influence of biological sex on MIF signalling within the uterus during pregnancy has not been shown before. Alternatively, VEGF signalling within the placenta has been shown to regulate vasculogenesis and angiogenesis processes and influence EVT maturation and invasion [56,57,71]. Surprisingly, we find male EVT, but not female EVT significantly engage PGF-FLT1 interactions with decidual endothelial cells. Further, macrophages in pregnancies with a male, but not a female fetus are predicted to engage in VEGF signalling with EVT and endothelial cells, defined in part by robust VEGF signalling components within these cell types [72]. While limited by sample size, our findings demonstrate that these signalling components may be differentially employed by maternal (female) immune cells found in the decidua depending on the corresponding sex of the placenta/trophoblast. Alternatively, AMOTL2 was found with increased expression in male HLA-G^+^ cells. AMOTL2 encodes a protein that has been shown to be involved in a variety of biological processes related to extravillous trophoblast differentiation and mature EVT function. This includes TGF-β inhibition via YAP1 phosphorylation [73], inhibition of the WNT signalling pathway in zebrafish [74], and the promotion of endothelial cell migration and proliferation during angiogenesis [75]. Further exploration of the relationship between VEGF signalling, angiogenesis, and increased AMOTL2 expression in male HLA-G^+^ cells is therefore needed to understand how placental sex influences vascular processes across the first trimester.

Finally, of the 20 autosomal transcripts found with increased expression in female HLA-G^+^ trophoblasts after week 10 of gestation, 7 are positively associated with preeclampsia (i.e., LOC101927354, PSG10P, PLEKHH1, LEP, MUC15, ANXA6, FAR2) [76–83] and 4 are negatively associated with preeclampsia (i.e., HS3ST3B1, ABCB1, MAFF, PKIB) [84–87]. Because pregnancies having a female fetus are suggested to be at higher risk of developing pre-eclampsia [21] future work is needed to explore the connection between the expression of these 11 genes in male versus female EVT and the associated onset/risk of preeclampsia.

### Perspectives and significance

Overall, our study resolves sex-dependent autosomal gene expression differences in human trophoblasts across the first trimester of pregnancy and identifies a key developmental window for the onset of sex-dependent autosomal gene expression. Importantly, these findings demonstrate that extravillous trophoblast plug erosion around week 10 of gestation correlates with a switch from a proliferative to immunomodulatory gene signature in extravillous lineage trophoblasts and results in the onset of distinct autosomal gene expression in male versus female HLA-G^+^ cells of the placenta. We further show that these sex-dependent autosomal gene expression differences influence how male and female EVT differentially employ the immunomodulatory MIF, CD99, and angiogenesis-associated VEGF signalling pathways across the first trimester uterine cell landscape.

## AUTHOR CONTRIBUTIONS

Conceptualization: M.J.S., A.G.B.; Methodology: M.J.S., A.G.B.; Formal analysis: bioinformatic – M.J.S., benchwork – M.J.S.; Resources: A.G.B.; Writing: original draft – M.J.S., review & editing – M.J.S., A.G.B.; Supervision: A.G.B.; project administration: A.G.B.; Funding acquisition: A.G.B.

## ACKNOWLEDGEMENTS

The authors extend their sincere gratitude to the hard work of staff at the British Columbia Women’s Hospital CARE Program for recruiting participants into our study and the staff at the University of British Columbia Biomedical Research Centre, especially Yiwei Zhao and Ryan Vander Werff, for assistance with bulk RNA-seq library sequencing. As well, we would like to thank Hoa Le for assisting with the collection of some of the HLA-G-enriched human trophoblast samples used in this study.

## ABBREVIATIONS

cCTB: Column cytotrophoblast
CTB: Cytotrophoblast
EVT: Extravillous trophoblast
hCG: Human chorionic gonadotrophin
HLA: Human leukocyte antigen
MC: Mesenchymal core
PCA: Principal component analysis
RNA-seq: RNA sequencing
SCT: Syncytiotrophoblast
SCTp: Syncytiotrophoblast precursor
UMAP: Uniform manifold approximation and projection

## TITLES AND LEGENDS TO SUPPLEMENTAL FILES

**Supplemental Figure 1:**
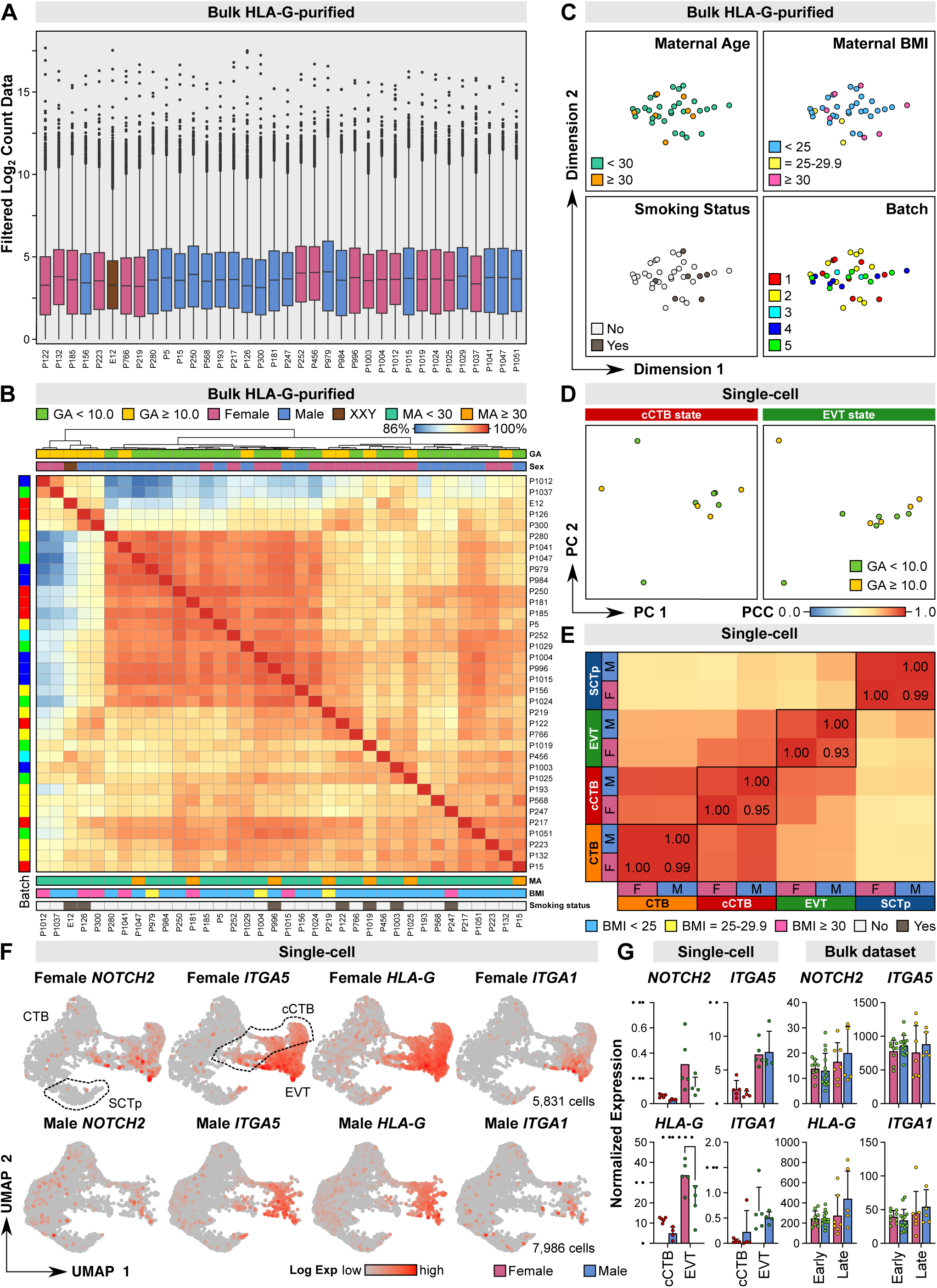
Bulk HLA-G^+^ trophoblast data quality control and verification against first trimester single-cell trophoblast data. (A) Box plots showing the filtered Log_2_ count data across all female, male, and XXY bulk HLA-G^+^ trophoblast samples. (B) Pearson’s correlation coefficient heatmap between all bulk RNA-seq samples. Sample metadata is indicated. Bulk RNA-seq Pearson’s correlation coefficient results are summarized in Table S2A. (C) Multidimensional scaling plots demonstrating sample clustering across principal component (PC) 1 vs. PC2. Samples are colour coordinated based on maternal age (years; top left), maternal body mass index (BMI, measured in kg/m^2^; top right), maternal smoking status (bottom left), or sequencing batch (bottom right). (D) Pseudo-bulk principal component analysis of cCTB state and EVT state samples colour-coded by gestational age category. (E) Pearson’s correlation coefficient heatmap between all female and male single-cell trophoblast states. Single-cell Pearson’s correlation coefficient results are summarized in Table S2B. (F) Expression levels (red) of known EVT-associated transcripts (NOTCH2, ITGA5, HLA-G, ITGA1) plotted in first trimester trophoblast single-cell UMAP space. (G) Bar plots demonstrating sample-specific gene expression levels of EVT-associated transcripts in the single-cell (left) cCTB and EVT states as well as the early and late first trimester male (XY) and female (XX) bulk RNA-seq (right) data (n=35) after removal of sample E12 (XXY).

**Supplemental Figure 2:**
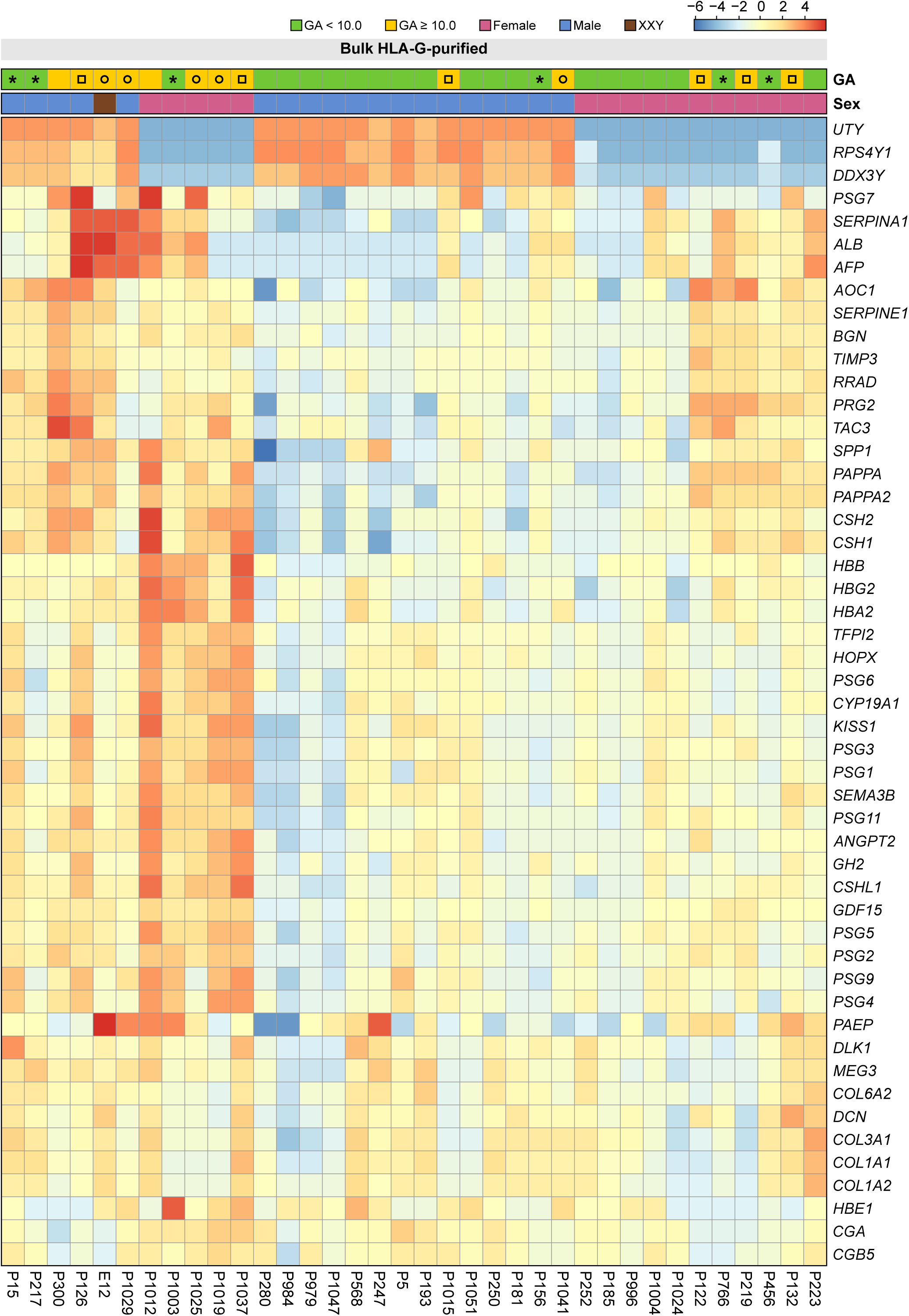
Heatmap of the top 50 most variably expressed transcripts identified in bulk HLA-G^+^ first trimester trophoblasts. Samples are clustered according to the dendrogram presented in Fig. 2A. Sample gestational age (GA) and sex are indicated. Asterisks indicate samples with a gestational age in week 9 of pregnancy, open circles indicate samples with a gestational age in week 10 of pregnancy, and open squares indicate samples with a gestational age in week 11 of pregnancy.

**Supplemental Figure 3:**
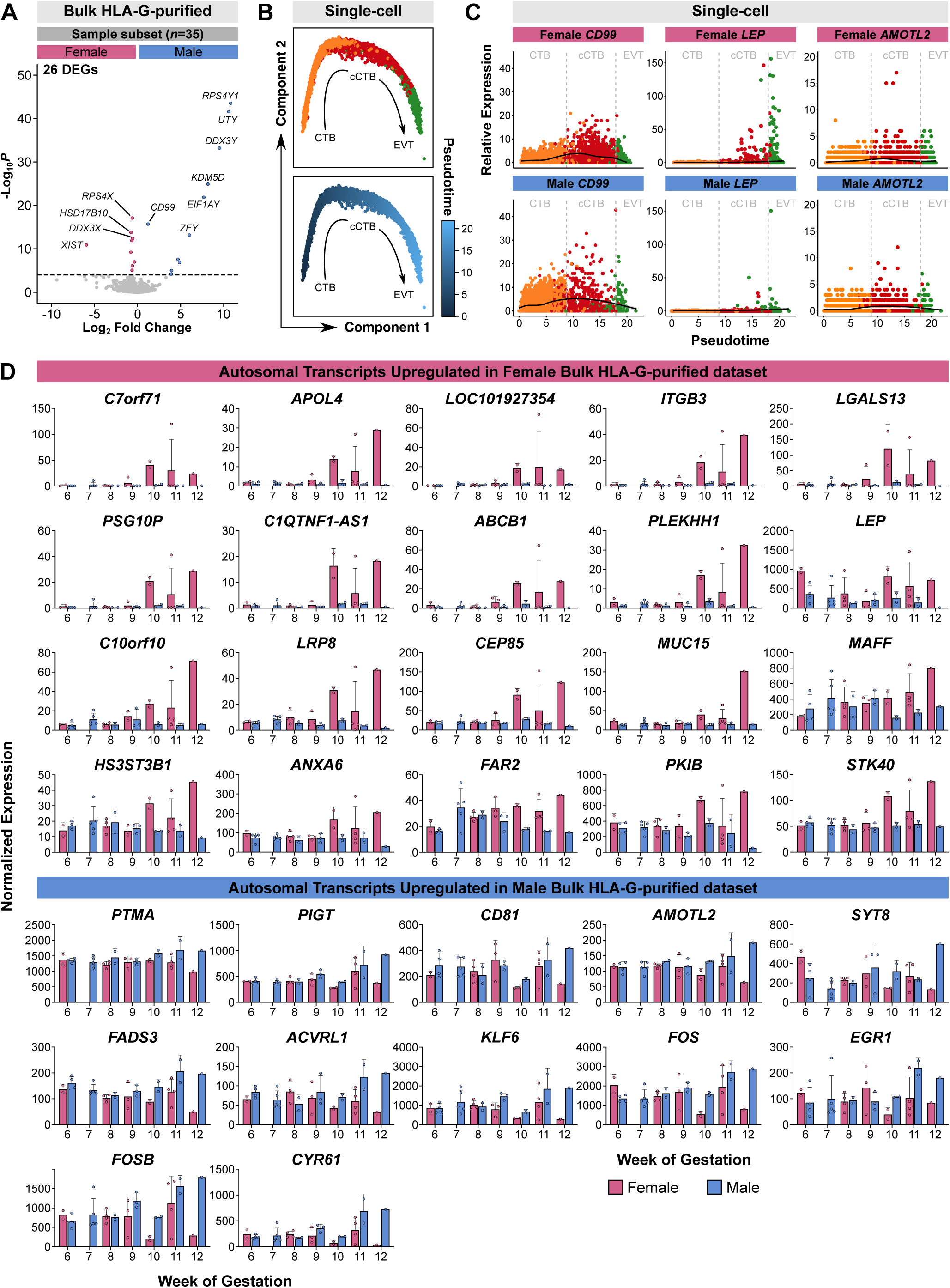
Autosomal sex differences arise around week 10 of gestation. (A) Volcano plot showing differentially expressed transcripts between all female and male first trimester HLA-G^+^ trophoblasts. (B) Monocle2 ordering of first trimester male and female single-cell CTB, cCTB, and EVT states colour-coded by cell state (top) or computed pseudotime value (bottom). (C) Dot plot showing female (top) and male (bottom) gene expression across predicted pseudotime value. Cells are colour-coded by cell state. (D) Bar plots demonstrating sample-specific gene expression levels of the significant autosomal transcripts identified in Fig. 3D. Samples are grouped into gestational week categories to observe the progression of sex-dependant female- and male-enriched autosomal gene expression across the first trimester.

**Supplemental Figure 4:**
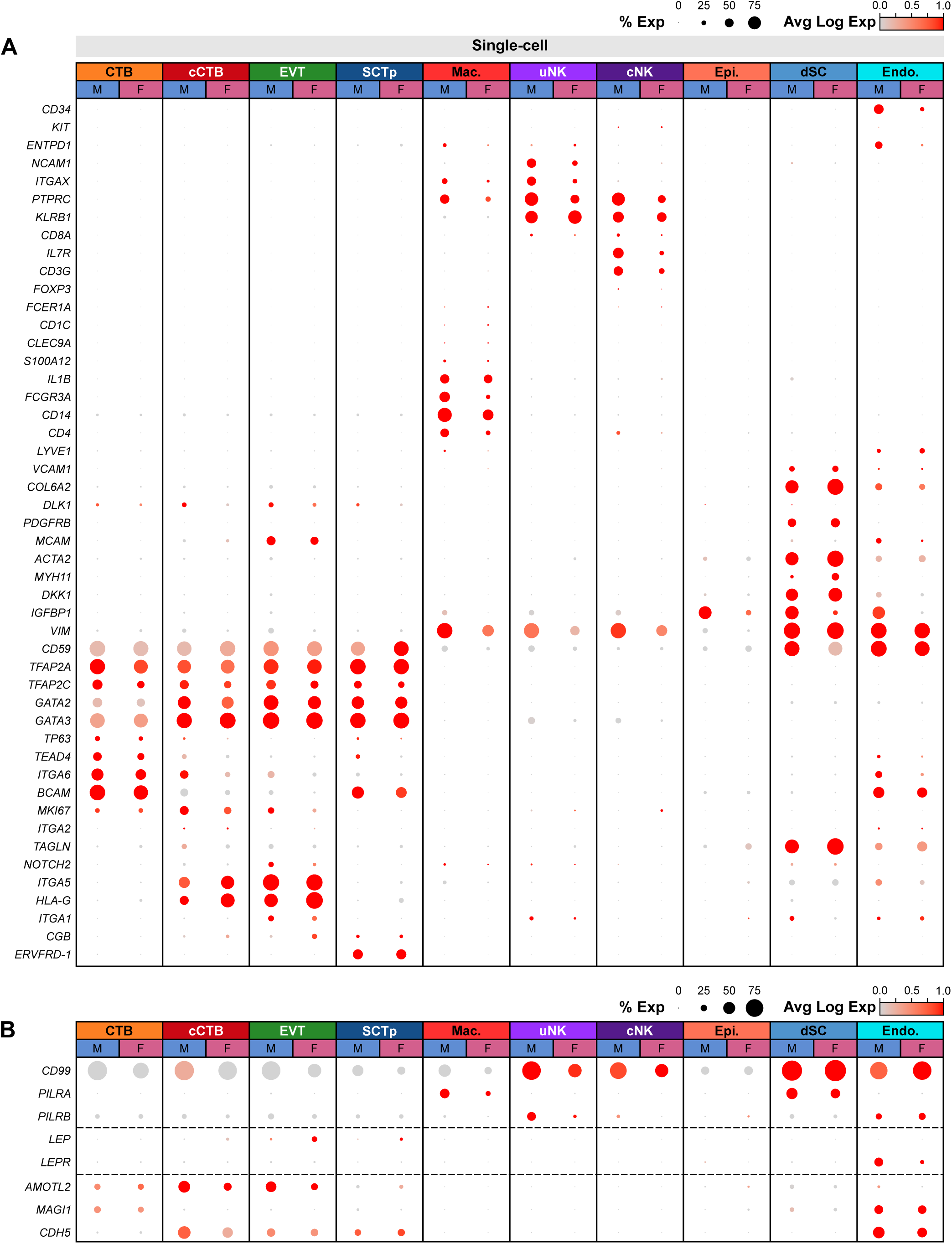
Maternal-placental interface cell state determination and gene expression. (A) Dot plot showing the percent and average Log expression level (0.0 to 1.0) of known trophoblast and decidual cell gene markers across all cell states in the expanded single-cell dataset. (B) Dot plot showing the percent and average Log expression level (0.0 to 1.0) of CD99, the CD99 receptors PILRA and PILRB, LEP and its receptor LEPR, as well as AMOTL2 and its binding partners MAGI1 and CDH5 in each cell state, split by sex.

